# 8-OxoG in GC-rich Sp1 binding sites enhances gene transcription during adipose tissue development in juvenile mice

**DOI:** 10.1101/538967

**Authors:** Jong Woo Park, Young In Han, Tae Min Kim, Su Cheong Yeom, Jaeku Kang, Joonghoon Park

**Author notes:** To whom correspondence should be addressed.; (J.P.). Correspondence may also be addressed to; (J.K.). The authors wish it to be known that, in their opinion, the first two authors should be regarded as joint First Authors.

## Abstract

The oxidation of guanine to 8-oxoguanine (8-oxoG) is the most common type of oxidative DNA lesion. There is a growing body of evidence indicating that 8-oxoG is not only pre-mutagenic, but also plays an essential role in modulating gene expression along with its cognate repair proteins. In this study, we investigated the relationship between 8-oxoG formed under intrinsic oxidative stress conditions and gene expression in adipose and lung tissues of juvenile mice. We observed that transcriptional activity and the number of active genes were significantly correlated with the distribution of 8-oxoG in gene promoter regions, as determined by reverse-phase liquid chromatography/mass spectrometry (RP-LC/MS), and 8-oxoG and RNA sequencing. Gene regulation by 8-oxoG was not associated with the degree of 8-oxoG formation. Instead, genes with GC-rich transcription factor binding sites in their promoters became more active with increasing 8-oxoG abundance as also demonstrated by specificity protein 1 (Sp1)- and estrogen response element (ERE)-luciferase assays in human embryonic kidney (HEK293T) cells. These results indicate that the occurrence of 8-oxoG in GC-rich Sp1 binding sites is important for gene regulation during adipose tissue development.

## INTRODUCTION

DNA, which carries genetic information, is continuously exposed to various types of damaging factors, including internally generated reactive oxygen species (ROS) along with external factors such as ionizing radiation, ultraviolet radiation, ROS, and other reactive molecules. These can cause DNA lesions that are processed by dedicated repair pathways to protect genome integrity. Unrepaired DNA lesions cause mutations that can lead to cell death or cancer (Ciccia and Elledge 2010).

ROS either from oxidative metabolism or exposure to agents that induce oxidative stress such as ionizing radiation or other environmental factors can result in oxidative DNA lesions (Olive and Johnston 1997). Over 100 different types of oxidative modification have been identified to date including damaged pyrimidines and purines as well as single strand breaks (SSBs) and abasic sites (Caldecott 2008). Guanine (G) has the lowest redox potential among nucleobases and is thus the most susceptible to oxidation; the oxidation of G to 8-oxoG (also known as 8-hydroxyguanine) is the most abundant and well-characterized oxidative DNA lesion whose repair is critical since it can pair with adenine (A) as well as cytosine (C) during replication, resulting in a G:C to T:A transversion mutation. The major repair pathway for these lesions is base excision-repair (BER) (Akiyama et al. 1989; Anson and Bohr 2000). In mammalian cells, 8-oxoG DNA glycosylase 1 (OGG1) recognizes and catalyzes the excision of 8-oxoG from the lesion, generating an abasic site. Apurinic/apyrimidinic endodeoxyribonuclease 1 (APEX1) cleaves the 5′ end of this site to generate an SSB with a 3′ hydroxyl and 5′ sugar phosphate. DNA polymerase β removes the latter through its lyase activity and fills the gap via templated DNA synthesis. DNA ligase I or III seals the nick and completes the BER process (David et al. 2007). Unrepaired 8-oxoG modifications have been implicated in cancer, neurodegenerative diseases, and aging (Radak and Boldogh 2010).

Genome-wide mapping of 8-oxoG in normal rat kidney has revealed that the lesion is preferentially located in gene deserts, and that there is no correlation between its distribution and gene expression (Yoshihara et al. 2014). *In situ* detection of 8-oxoG using a monoclonal antibody on a human metaphase spread prepared from peripheral lymphocytes revealed that 8-oxoG is non-uniformly distributed in the normal human genome and that the pattern is conserved across individuals. Additionally, a high density of 8-oxoG coincides with regions exhibiting a high frequency of recombination and single nucleotide polymorphisms (Ohno et al. 2006).

There is increasing evidence of an association between DNA repair and transcriptional regulation. Oxidized G lesions function in OGG1-mediated epigenetic regulation of nuclear factor-κ B (NFκB)-induced gene expression, and OGG1 enhances pro-inflammatory gene expression by facilitating the recruitment of site-specific transcription factors (Pan et al. 2016). On the other hand, 8-oxoG may act as on/off switch for transcription depending on the strand on which it is located in the context of a gene promoter G-quadruplex (Fleming et al. 2017b).

In this study we carried out genome-wide 8-oxoG profiling of adipose and lung tissues of juvenile female C57BL/6 mice by affinity purification followed by next-generation sequencing in order to clarify the genetic and molecular roles of 8-oxoG beyond its function as a DNA damage mark. We found that transcriptional activity and the number of active genes were correlated with 8-oxoG distribution, especially in gene promoters. A transcription factor binding motif analysis revealed that genes that were highly expressed - especially in adipose tissue - had GC-rich promoters as compared to those that were moderately active or inactive genes. Furthermore, genes with GC-rich transcription factor binding sites in their promoters became more active with increasing 8-oxoG abundance as demonstrated by Sp1- and ERE-luciferase assays in HEK293T cells under oxidative stress condition. These results suggest that 8-oxoG promotes transcription during adipose tissue development in mice.

## RESULTS

### 8-oxoG is abundant in the genome of various tissues of juvenile mice

Hydrolyzed genomic DNA samples from each tissue were analyzed by RP-LC/MS to determine 8-oxoG levels. For quality assurance of the procedure, we also measured total dG and dC by HPLC. Representative chromatograms and standard curves generated with various concentrations of 8-oxoG standard are shown in Supplementary Figure S1. The retention time of 8-oxoG was 2.9 min, and the correlation coefficient (*R*^2^) was 0.999 with 8-oxoG standard. Averaged 8-oxoG levels with standard deviation (SD) in lung, liver, and adipose tissues were 0.025% ± 0.008%, 0.054% ± 0.012%, and 0.070% ± 0.008% of total dG, respectively (Table 1). Coefficient of variation of 8-oxoG levels in different tissues was less than 32%. These global concentrations of 8-oxoG were comparable to the background range of 8-oxoG levels in pig liver (0.0002% to 0.0441%), HeLa cells (0.0001% to 0.0214%) (Escodd 2002) or commercially available calf thymus DNA (0.032%) (Chepelev et al. 2015). The ratios of total dG to total dC were 1.03, 1.00, and 0.91 in lung, liver, and adipose tissues, respectively, indicating that there was no detection bias in the analysis. The level of 8-oxoG was lower in lung tissues than in liver or adipose tissues (*P* < 0.01). These results demonstrate that 8-oxoG is present at a low but significantly different level in genomic DNA of adipose and lung tissues of juvenile mice, and these tissues were subjected to further analyses. Since we used sexually immature mice, we might be able to avoid the effects of aging on 8-oxoG formation.

**Table 1.** Global levels of 8-oxoG in various tissues of juvenile female mice

| Tissue type | No. of animals | %8-oxoG/dG<sub>total</sub> | dG<sub>total</sub>/dC<sub>total</sub> |
| --- | --- | --- | --- |
| Lung | 5 | 0.025 ± 0.008 | 1.032 ± 0.018 |
| Liver | 4 | 0.054 ± 0.012** | 1.005 ± 0.051 |
| Adipose | 4 | 0.070 ± 0.008*** | 0.910 ± 0.088 |

Mean ± Standard Deviation (SD), \*\* $P < 0.01$ , \*\*\* $P < 0.001$ vs. lung, one-sided Student's $t$ -test.

### Genomic distribution of 8-oxoG and effects on gene expression

We performed OG-seq to determine the genomic location of 8-oxoG lesion as well as RNA sequencing to clarify the association between 8-oxoG and gene activity. More than 120 million mappable reads - of which 95.4% had a lower base call accuracy of 99% (Q20) by OG-seq - were generated for each tissue. An average of 32,284 and 70,848 8-oxoG peaks with significant fold enrichment (*Q* < 0.01) compared to the input DNA were identified in adipose and lung tissues, respectively (Supplementary Table S1 and S2). The chromosomal locations of the enriched 8-oxoG peaks were annotated in terms of gene symbols and genetic elements. Additionally, a median of 118 million mappable reads with 94.3% Q20 generated by RNA sequencing were annotated; their expression levels are summarized in Supplementary Table S3 as FPKM values. We integrated the enriched 8-oxoG peaks and gene expression levels in a Circos plot, which revealed that 8-oxoG lesions as well as the active genes were evenly distributed throughout the chromosomes of each tissue (Figure 1 and Supplementary Figure S2). Although the total number of 8-oxoG peaks was higher in lung than in adipose tissue, the opposite was true for the concentration of 8-oxoG peaks in a specific region, which is consistent with the global levels of 8-oxoG lesions determined by LC/MS in each tissue.

**Figure 1.**
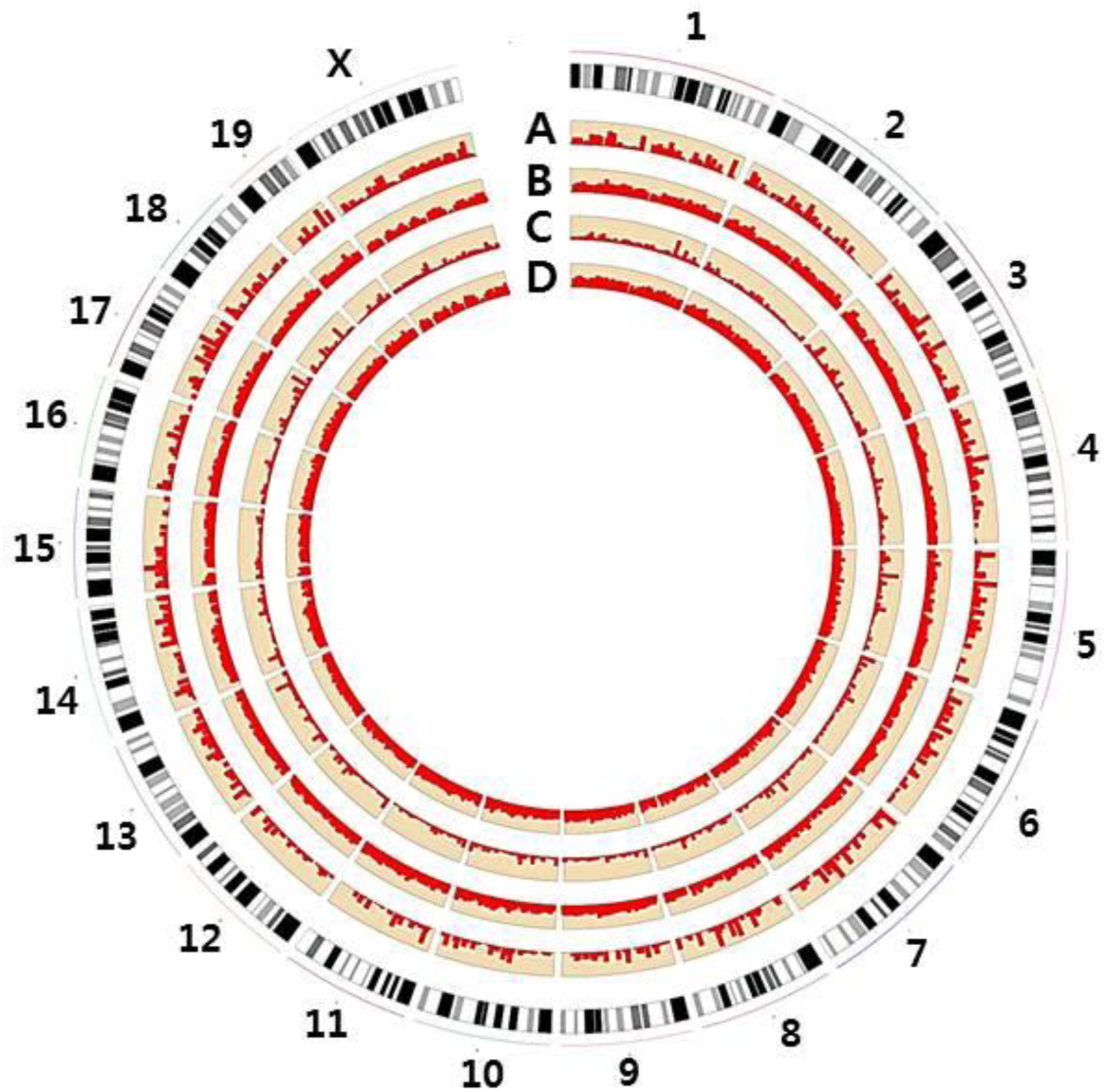
Integration of genomic distribution of 8-oxoG and gene expression in adipose and lung tissues of juvenile mice. In the Circos plot, the outer layer indicates the juvenile female mouse genome (mm10); the inner layers depict tissue-specific 8-oxoG distribution and gene expression. (**A, C**) Genome-wide distribution of significantly enriched 8-oxoG peaks (*Q* < 0.01) and (**B, D**) global gene expression levels in (**A, B**) adipose and (**C, D**) lung tissues. The degree of 8-oxoG concentration ranged from 10 to 40, and gene expression levels were log transformed (log_10_).

We then identified expressed genes harboring 8-oxoG lesions in their DNA sequence. Among the 18,963 genes commonly expressed in both tissues, 778 genes in adipose and 584 genes in lung tissue had tissue-specific 8-oxoG formation. By contrast, 1,472 genes had 8-oxoGs with non-tissue-specific manner (Figure 2A). There were 436 and 398 genes in adipose and lung tissue, respectively, those were not expressed but had tissue-specific 8-oxoG lesions. By contrast, 1,466 genes not expressed in both tissues had 8-oxoGs in common. Genes with 8-oxoGs were classified as high, low, or off genes depending on their expression level. The number of genes designated as off, low, and high genes was 436, 457 and 321, respectively, in adipose tissue (Figure 2B) and 398, 283, and 301, respectively, in lung tissue (Figure 2C).

**Figure 2.**
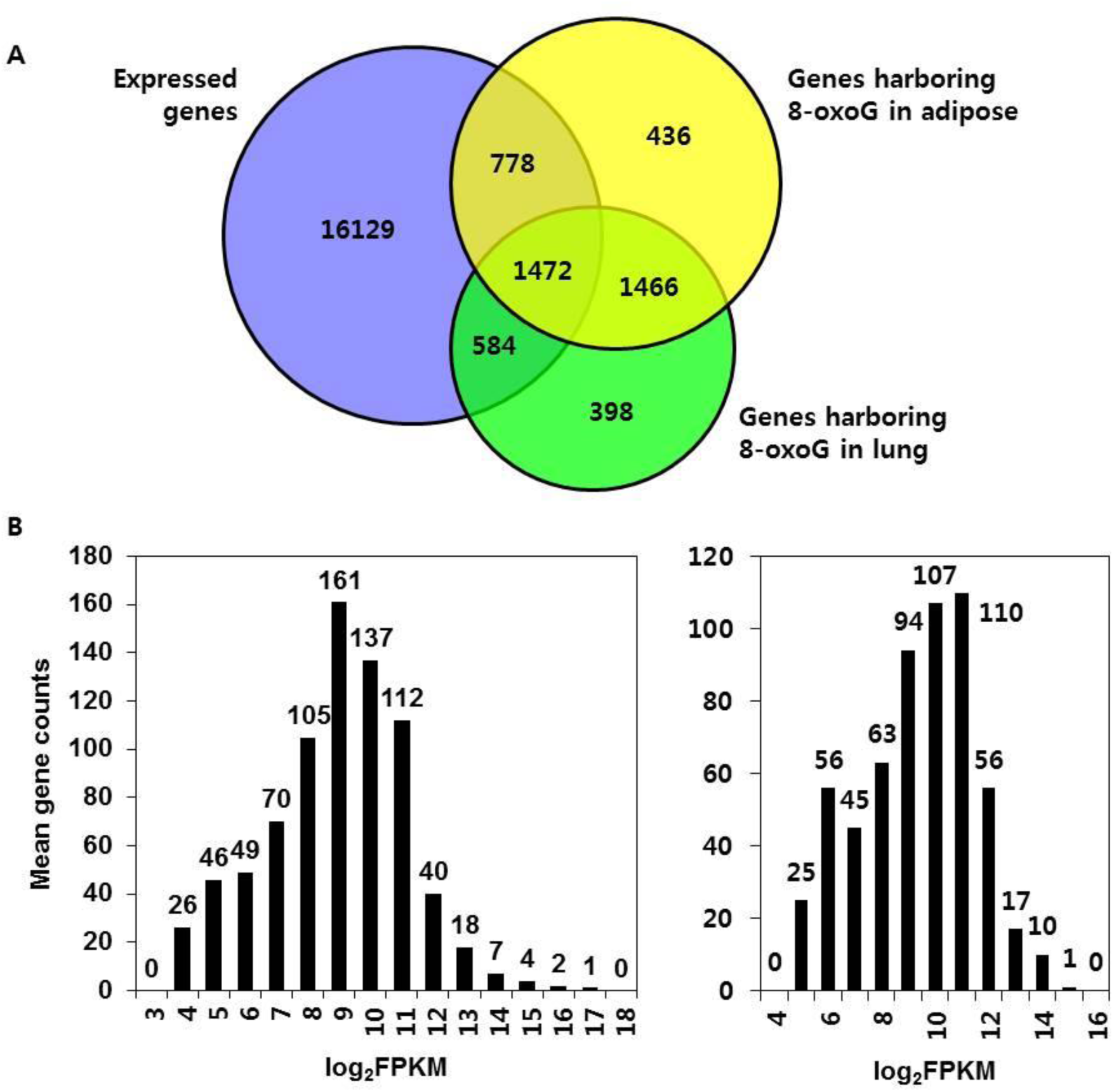
Correlation analysis between gene activity and 8-oxoG abundance. (**A**) Venn diagram of commonly genes expressed in both of adipose and lung tissues (purple), genes with 8-oxoG lesions in adipose tissue (yellow) and lung tissue (green). Histograms showing the distribution of gene counts according to expression level in (**B**) adipose and (**C**) lung tissues. Gene expression levels were log transformed to FPKM values.

The location of 8-oxoG peaks in each tissue was classified as 3’ untranslated region (UTR), 5’ UTR, distal intergenic, downstream, exon, intron, or promoter region. When we compared the fraction of 8-oxoG peaks in each genetic element according to gene activity, we found that peaks were less abundant in the distal intergenic regions and more prevalent in promoter regions in association with increased gene expression levels. In adipose tissue, averaged 63.2% of 8-oxoG peaks in intergenic regions were in off genes, 41.8% in low genes, and 33.9% in high genes (Figure 3A). In contrast, 13.2% of peaks were located in the promoter region of off genes, 26.7% in that of low genes, and 42.5% in that of high genes (*P* < 0.01, R^2^ = 0.9394). A similar 8-oxoG distribution was observed in lung tissues (Figure 3B). The averaged proportion of the 8-oxoG peaks in promoter regions was 27.1% for high genes, which was significantly higher than the proportion of 8-oxoG peaks in the promoter of off and low genes (19% and 20.9%, respectively; *P* < 0.01, R^2^ = 0.3668). Read count frequency analysis revealed that the peaks were localized around gene transcription start site (TSS) (Supplementary Figure S3 and S4).

**Figure 3.**
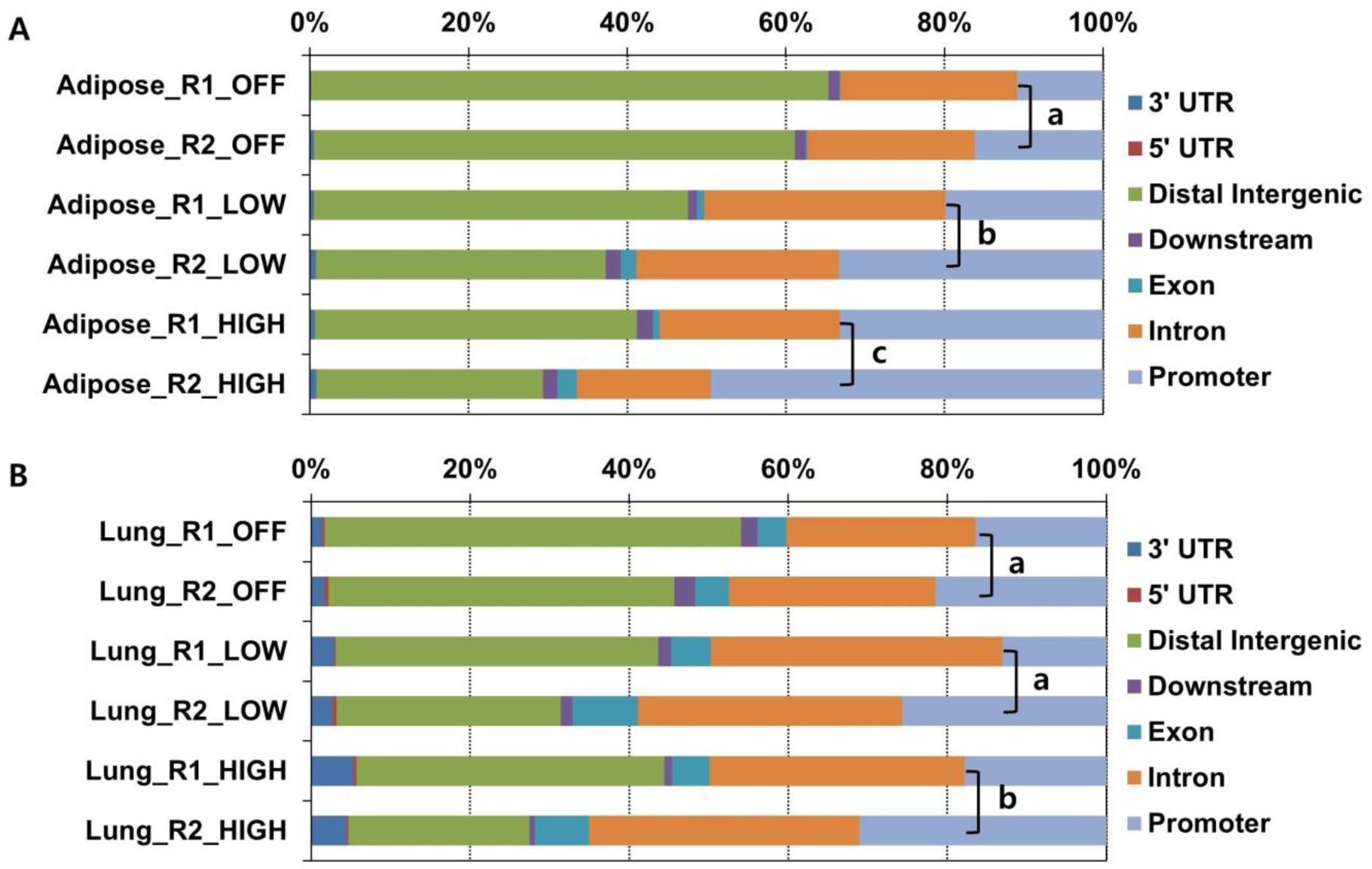
Genomic distribution of 8-oxoG in adipose and lung tissues. Proportions of genomic distribution of 8-oxoG in (**A**) adipose and in (**B**) lung tissues. X axis denotes the relative proportion of 8-oxoG in each genetic element, Y axis denotes the categorized genes according to expression levels. Different lowercase letters in the promoter region indicate significant differences with Fisher’s exact test (*P* < 0.01 vs. a).

The observed density of 8-oxoG in the gene promoters implied that these lesions are related to gene activity. One possibility is that promoters have an abundance of 8-oxoG to increase gene activity. Alternatively, promoter regions may have a unique DNA sequence context that employs 8-oxoG to regulate gene activity in an epigenetic fashion. To test the first hypothesis, we normalized the fold enrichment and the number of 8-oxoG peaks according to the promoter length of each gene, and then evaluated the correlation with gene expression. The 8-oxoG peaks were enriched 1.8 fold in off genes, 1.7 fold in low genes, and 1.7 fold in high genes in the promoter regions without a significant difference among gene categories (Figure 4A). Likewise, there were no differences in the number of the 8-oxoG peaks per 100-bp promoter length (Figure 4B; 0.57, 0.55, and 0.56 8-oxoG peaks per 100-bp promoter length for off, low, and high genes, respectively). These results imply that 8-oxoG-associated gene regulation is unrelated to 8-oxoG abundance in the promoter regions.

**Figure 4.**
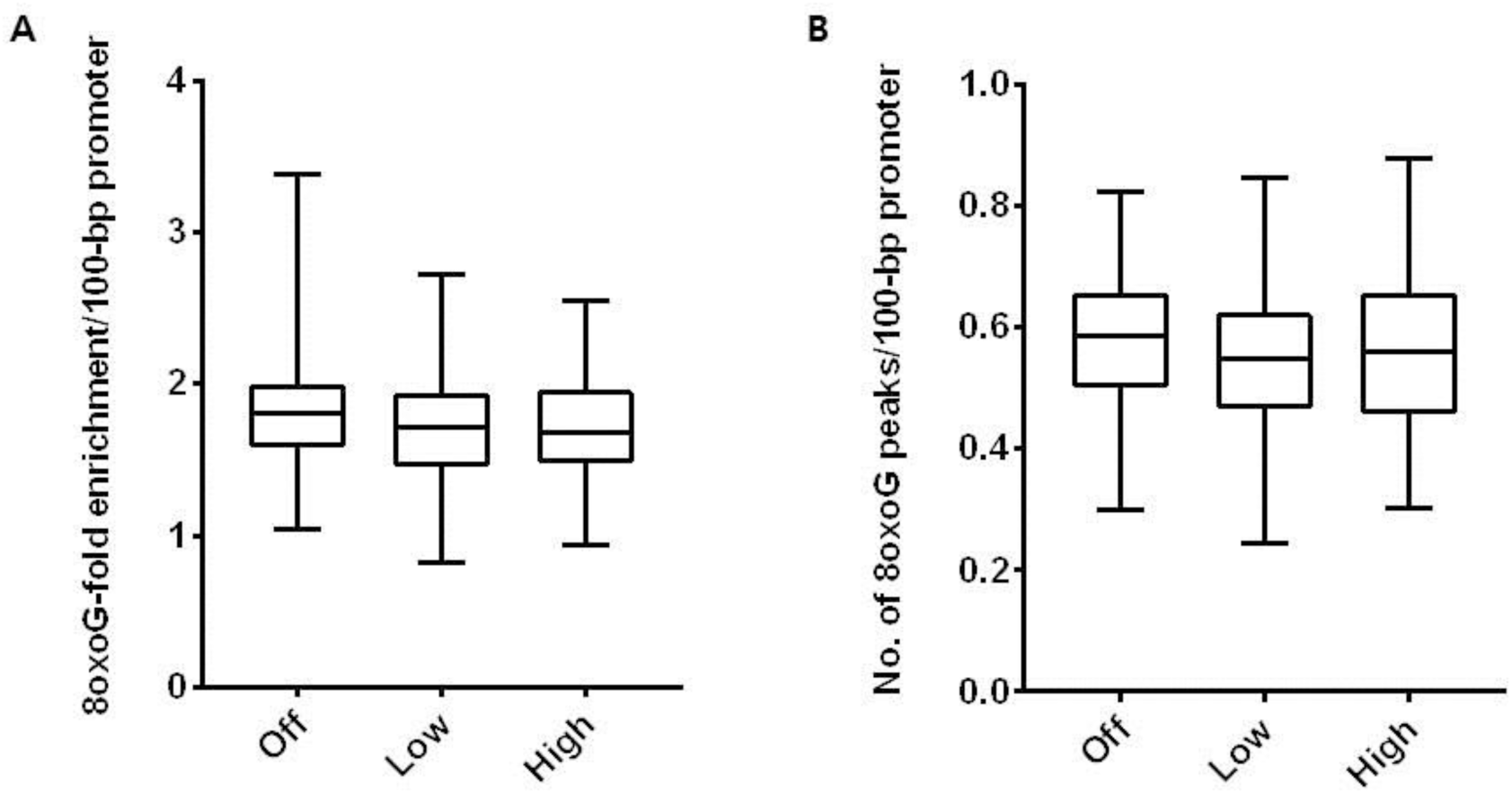
Correlation between gene expression and 8-oxoG enrichment in promoter regions in adipose tissue. Fold enrichment of (**A**) 8-oxoG and (**B**) number of 8-oxoG peaks per 100-bp promoter length compared to input DNA. Genes were categorized according to expression level.

### Transcriptional binding motif of genes with promoters harboring 8-oxoG in adipose tissue

Next, we investigated the feature of the DNA sequence of the promoter regions harboring 8-oxoG lesions to clarify the significance of 8-oxoG-associated gene regulation. Based on the observed association between gene activity and higher 8-oxoG abundance in promoter regions in adipose tissues, we performed functional enrichment analyses of adipose tissue-specific genes. We found that off genes with 8-oxoGs were functionally enriched in apoptotic process (*P* = 4.87E-06) and cell death (*P* = 5.76E-06), whereas low or high genes with 8-oxoGs were enriched in regulation of gene expression (*P* = 8.36E-13), cell cycle (*P* = 1.01E-08), sequence-specific DNA binding (*P* = 5.79E-09), and catabolic process (*P* = 3.36E-08) (Supplementary Figure S5). These results demonstrate that as expected, genes required for the survival and metabolic function of adipose tissue are active whereas those involved in cell death are silenced in juvenile mice.

The regulation of adipose tissue-specific gene activity in these mice was closely related to the occurrence of 8-oxoG lesions. To examine DNA context-dependent gene regulation by 8-oxoG in greater detail, we performed an enrichment analysis of transcription factor binding motifs for the genes and found that those with GC-rich transcription factor binding sites in their promoters became more active as a function of 8-oxoG abundance. While only four off genes were enriched in the GC-rich Sp1 binding motif (Figure 5A; *P* = 2.44E-05), 31 low genes (Figure 5B; *P* ≤ 1.44E-09) and 55 high genes (Figure 5C; *P* ≤ 3.92E-16) in adipose tissue harbored a large number of GC-rich Sp1, Paired box 4 (Pax4), or Myc-associated zinc finger protein (Maz) binding sites (Supplementary Tables S4-S6). In contrast, there was no enrichment of GC-rich transcription factor binding motifs according to gene activity in the promoter regions in genomic DNA from lung tissue (Supplementary Figure S6).

**Figure 5.**
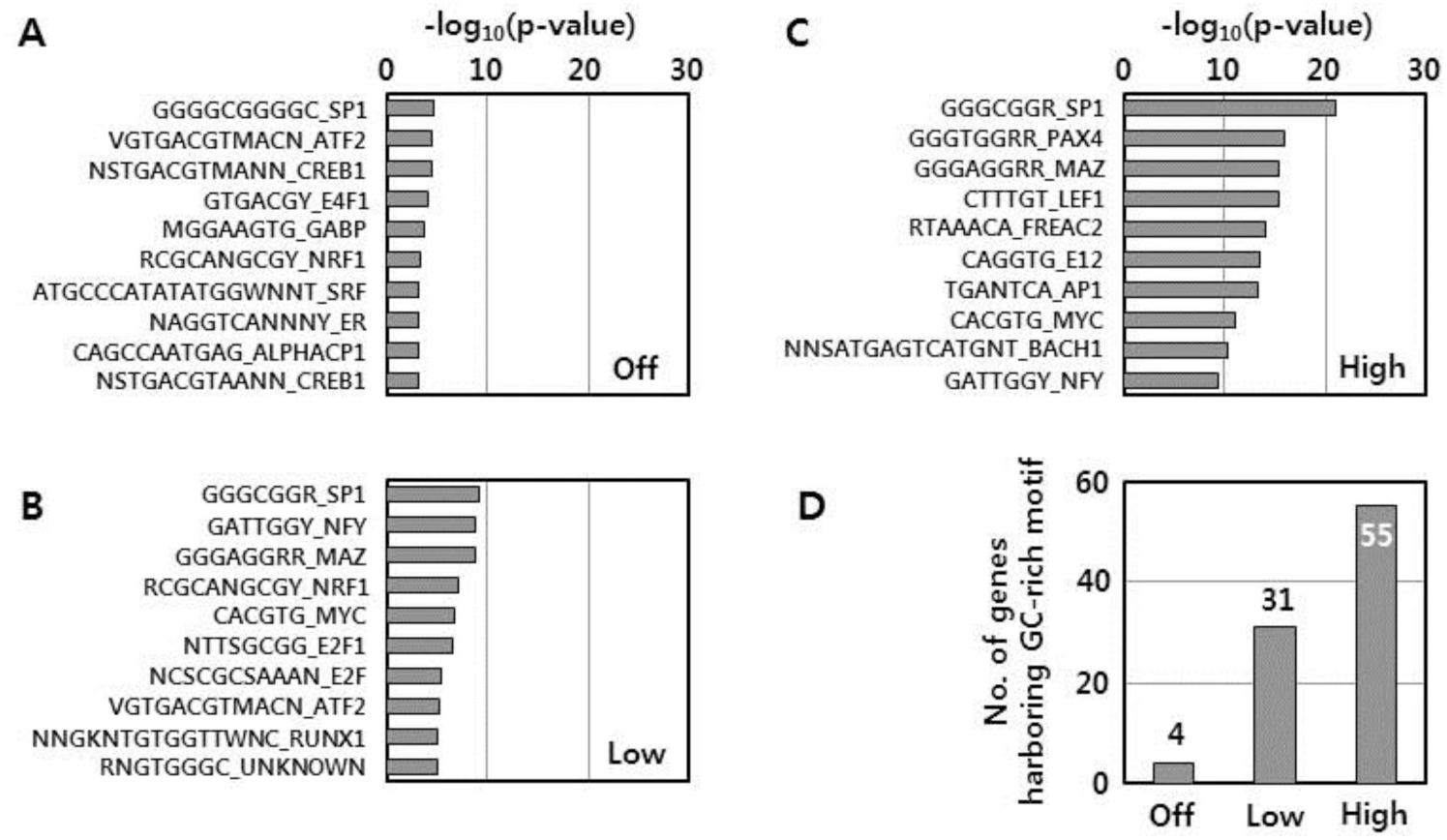
Transcription binding motif of genes with a promoter harboring 8-oxoG in adipose tissue. Enrichment analysis of transcription factor binding motifs of (**A**) off genes, (**B**) low genes, and (**C**) high genes. Enrichment P values were determined after log transformation. (**D**) Number of genes with GC-rich transcription factor binding sites such including *Sp1, Pax4*, and *Maz* according to gene expression level.

We carried out a literature search to clarify the biological relevance of genes harboring GC-rich transcription factor binding motifs with 8-oxoG lesions in adipose tissue. Among the 121 high genes, 47 genes (38.8%) were related to adipose tissue physiology according to at least one report, and 31 genes (66.0%) were predicted to have GC-rich transcription factor binding sites in the promoter (Supplementary Table S7). For example, Patatin-like phospholipase domain-containing 2 (*Pnpla2*) is an adipose triglyceride lipase that regulates lipid metabolism in adipose tissue (Lord and Brown 2012; Grahn et al. 2014; Dettlaff-Pokora et al. 2016). A genomic search revealed that there were five 8-oxoG peaks within the 3 kb up- or downstream of the *Pnpla2* gene TSS in adipose tissues, all of which were located in GC-rich Sp1 binding sites (Figure 6). Likewise, Nuclear receptor subfamily 1 group D member 1 (*Nr1d1*) (Fontaine et al. 2003; Laitinen et al. 2005), Cluster of differentiation 68 (*Cd68*) (Pietilainen et al. 2006; Moreno-Navarrete et al. 2013) and Sp1 (Prieto-Hontoria et al. 2011; Roy et al. 2017) - which encode regulators of adipocyte differentiation and metabolic function - had several 8-oxoG peaks with GC-rich Sp1 binding sites around the TSS. In contrast to the high genes, only 29 genes of 95 low genes (30.5%) and 14 genes of 50 off genes (28.0%) were demonstrated in at least one study to be related to adipose physiology. Only two of the off genes had GC-rich transcription factor binding motifs, which was significantly lower than for active genes (*P* < 0.0154). These results indicate that genes essential for the development and function of adipose tissue are upregulated by 8-oxoG lesions and that adipose tissue-specific genes have GC-rich promoters that are regulated by the epigenetic function of 8-oxoG.

**Figure 6.**
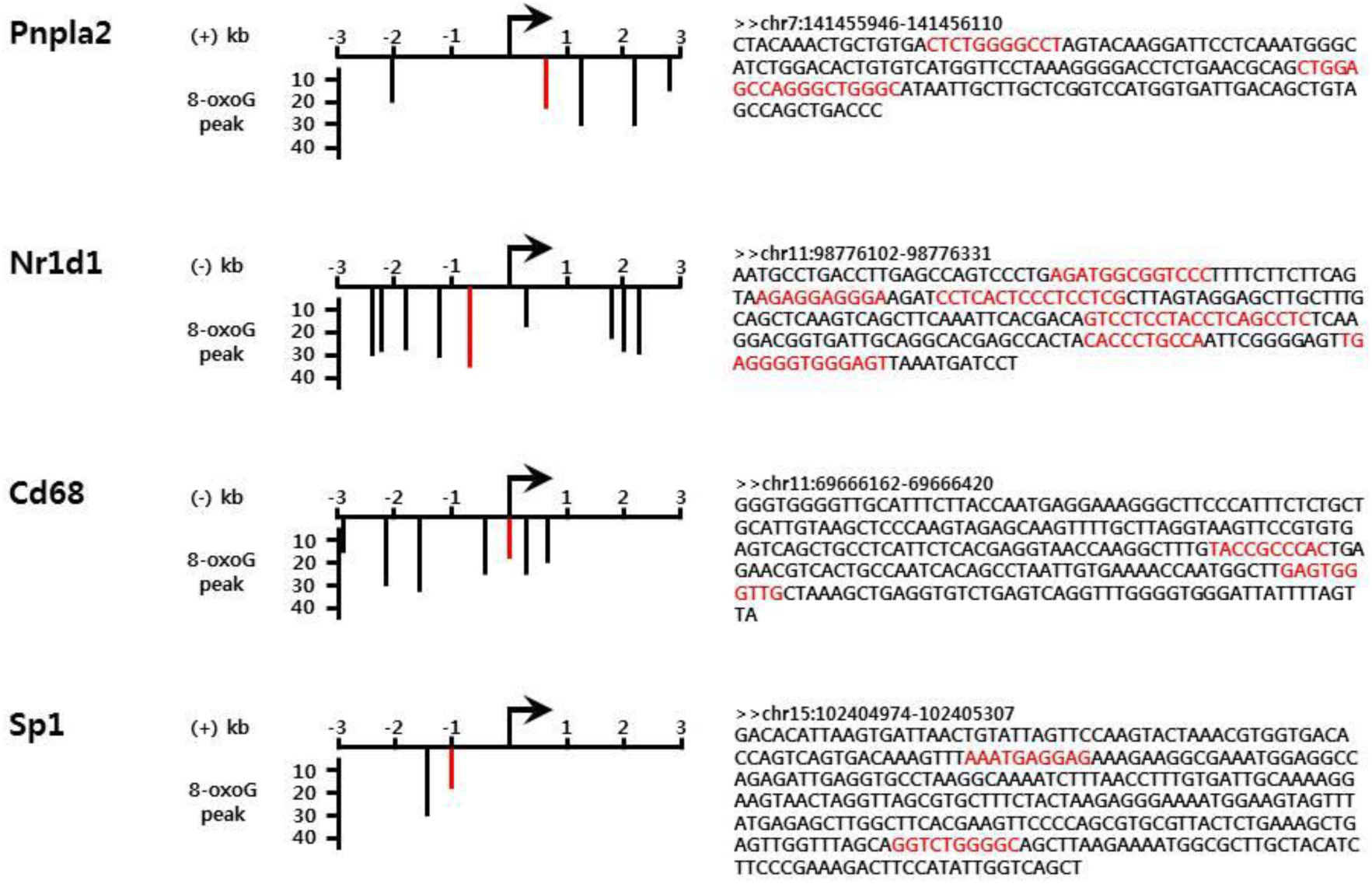
Genomic browser views of 8-oxoG peaks in the promoters of genes highly expressed in adipose tissue. Representative adipose tissue-specific functional genes with 8-oxoG concentrated around the TSS are shown. Gene direction is indicated by the (+) or (-) strand. Red bars represent 8-oxoG peaks and their DNA sequence shown to the right; Sp1 binding sequences were highlighted in red.

### GC-rich transcription factor binding motif-dependent gene activation

To verify whether the presence of 8-oxoG lesions is associated with GC-rich transcription binding site-dependent gene activation, we carried out luciferase assays using HEK293T cells. Sp-1 luciferase vector that we transfected into the cells contains high GC contents in the promoter region while ERE-luciferase vector contains low GC contents. First we tested whether treatment of the H_2_O_2_ to HEK293T cell could induce 8-oxoG lesion and it could be inhibited by pre-treatment of NAC. We observed immunofluorescently that 8-oxoG was induced by 300 µM H_2_O_2_ treatment and it was inhibited by co-treatment of 500 µM NAC (Figure 7A). There were no significant cell death upon the concentration of H_2_O_2_ or NAC used either standalone or combinational treatment (Figure 7B). When cells were subjected to oxidative stress by treatment with 300 µM H_2_O_2_, luciferase activity was significantly increased 5.4-fold in Sp1-transfected cells compared to the control group (*P* < 0.01). This effect was abrogated by co-administration of 500 µM NAC (Figure 7C). In contrast, oxidative stress-induced gene activation was not observed in ERE-transfected cells (Figure 7D). Taken together, gene activation in response to oxidative stress appears to be dependent on the DNA context of the transcription factor binding site, and GC-rich Sp1 binding sites would promote gene activation under the oxidative stress conditions.

**Figure 7.**
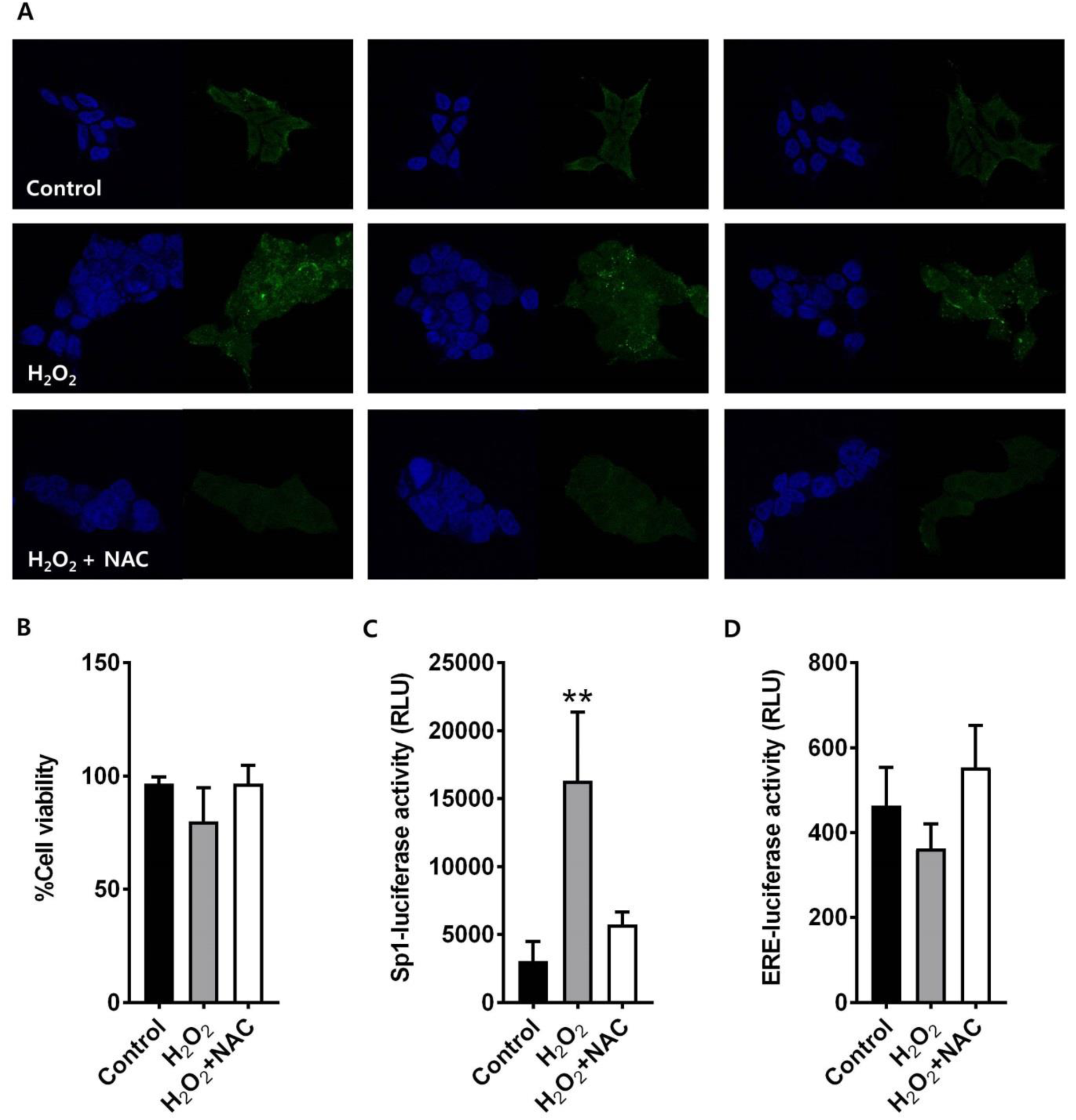
Immunofluorescence for 8-oxoG formation and inhibition upon H_2_O_2_ and NAC with DAPI counterstaining staining in HEK293T cells. (**A**) 8-OxoG formation by extrinsic H_2_O_2_ and/or NAC treatment. (**B**) Cell survival assays in response H_2_O_2_ either in presence or absence of NAC. (**C**) Sp1 site-mediated transcriptional activation under oxidative stress. (**D**) ERE site-mediated transcriptional activation under oxidative stress. Mean differences between groups were evaluated by analysis of variance followed by Dunnett’s multiple comparison test. \*\**P* < 0.01 vs. control.

## DISCUSSION

ROS formed as a result of oxidative stress are electron-deficient and readily oxidize proteins, lipids, RNA, and especially DNA (Fleming and Burrows 2017). Of the four DNA bases, heterocyclic G is the most susceptible to oxidation, with 8-oxoG as the major oxidatively modified product (Cadet et al. 2014). The long-standing view is that 8-oxoG is mutagenic and detrimental to cellular processes such as gene transcription. However, there is increasing evidence that in addition to being a pre-mutagenic DNA lesion, 8-oxoG also plays an essential role in the regulation of gene expression along with the DNA repair protein OGG1 (Wang et al. 2018).

8-OxoG formation in promoters followed by OGG1 recruitment may induce NF-κB-driven pro-inflammatory gene expression (Pan et al. 2016). It was also demonstrated that sirtuin 1 (*SIRT1*) gene expression is regulated by apurinic/apyrimidinic endonuclease 1 - a multifunctional protein contributing to BER - through oxidation of G to 8-oxoG in negative calcium responsive elements in the *SIRT1* promoter in HeLa cells under conditions of oxidative stress (Antoniali et al. 2014). This finding implies that both the abasic site and BER enzyme are key factors for gene activation associated with 8-oxoG formation in gene promoters. It was also reported that 8-oxoG formation in a putative G-quadruplex-forming sequence in the coding strand of the vascular endothelial growth factor (*VEGF*) promoter increased gene transcription (Fleming et al. 2017a). On the other hand, 8-oxoG in template strands was shown to block the advancement of RNA polymerase II and recruit the transcription-coupled repair machinery, thereby slowing transcription (Tornaletti et al. 2004; Guo et al. 2013). 8-OxoG was also found to inhibit transcription when located in either the coding or template strand in a gene-coding region (Kitsera et al. 2011; Allgayer et al. 2016).

Despite the controversy surrounding whether 8-oxoG positively or negatively regulates gene activity, the above studies suggest that 8-oxoG is an important epigenetic regulator of gene expression. However, most studies have focused on single genes under specific conditions, and it remains unclear whether 8-oxoG lesions are randomly formed in the genome or whether oxidation occurs in a sequence-specific manner. Thus, a major challenge for studying the role of 8-oxoG in gene regulation is the development and implementation of genome-wide 8-oxoG sequencing. In this study, we investigated the association between 8-oxoG formed in response to intrinsic oxidative stress and gene expression. The genome-wide analysis of 8-oxoG was performed with both adipose and lung tissues, which are representative tissues as insensitive and sensitive to oxygen toxicity, respectively. We also used juvenile mice to minimize the effects of aging on 8-oxoG formation.

Many laboratories have developed antibody-based 8-oxoG sequencing methods that provide a low-resolution sequence map of 8-oxoG distribution (approximately 10-1000 kb) (Ohno et al. 2006; Yoshihara et al. 2014). However, these do not supply information on the sequence-specific distribution of 8-oxoG and its epigenetic functions. In this study, we examined the genome-wide 8-oxoG distribution by applying OG-Seq at ~150-bp resolution using a chemical method to label 8-oxoG with biotin for affinity purification and sample enrichment (Ding et al. 2017). Through this advanced approach, we found a correlation between 8-oxoG distribution and gene expression. Transcriptional activity and the number of active genes were significantly correlated with the distribution of 8-oxoG, especially in the gene promoter regions. However, the 8-oxoG peaks and number of 8-oxoG peaks per 100-bp promoter length did not differ among genes categorized as inactive or moderately or high active. Instead, genes with GC-rich transcription factor binding sites in their promoters became more active with increasing 8-oxoG abundance, implying that 8-oxoG-related gene regulation is not associated with the degree of 8-oxoG formation but with DNA context within the promoter. The importance of 8-oxoG formation for GC-rich transcriptional binding site-dependent gene activation was confirmed by Sp1- and ERE-luciferase assays in HEK293T cells. Thus, oxidative stress-mediated gene activation is dependent on the DNA motif of the transcription factor binding site, with GC-rich Sp1 binding sites being a key element in the modulation of gene activity under oxidative stress.

Guanine oxidation does not occur randomly in the genome, but instead shows a strong distributional bias (Ba and Boldogh 2018). Furthermore, 8-oxoG substitution in synthetic DNA oligomer affected the binding affinity of Sp1 depending on guanine position (Ramon et al. 1999). However, the mechanisms by which 8-oxoG formation at GC-rich transcription factor binding sites and regulation of gene expression remain unclear. It has been proposed that in response to an oxidative burst, guanine oxidation in a GC-rich promoter has a *cis* effect, whereas OGG1 acts as a *trans* factor whose oxidation is deemed to be a reversible post-translational modification (Ba and Boldogh 2018). The rapid binding of oxidatively inactivated OGG1 and consequent allosteric alteration of DNA during the pre-excision step of BER (Koval et al. 2004) facilitate the homing of transcription factors (e.g., NF-κB and signal transducer and activator of transcription 1) or co-activator (e.g., C-terminal binding protein/p300) to their cognate binding sites, which promotes assembly of the transcriptional machinery. When the redox balance is re-established, OGG1 regains its enzymatic activity through reduction of its oxidized cysteine and the damaged base is then excised to prevent mutation(s) in the promoter.

There are some potential pitfalls to consider in this study. First, number of mice used in comprehensive genomic analysis of 8-oxoG was low (n = 2 per each tissues) with limited variety of tissues (lung and adipose tissues). Although we observed the significant enrichment of 8-oxoG in the GC-rich transcription factor binding motif-containing promoter regions of adipose tissue-specific genes according to the gene activity, there was no enrichment like this in genomic DNA from lung tissues. It turns to be elusive if the conclusion drawn in this study is general for other species and tissues. Second, the effect of glycosylase Ogg1 on the formation and distribution of 8-oxoG in genomic DNA was not investigated. We observed significantly reduced level of 8-oxoG in the lung tissues compared to the adipose tissues, which was well explained by differential expression of OGG1 in human lung and adipose tissues (https://www.proteinatlas.org/ENSG00000114026-OGG1/tissue). However, the precise role of Ogg1 in 8-oxoG formation and the transcriptional regulatory activity were beyond the scope of this study, and were remained to be clarified. Third, there would be difference in oxidative stress conditions between *in vivo* and *in vitro* experiments. We applied intrinsic oxidative stress condition in animal experiment and extrinsic oxidation using H_2_O_2_ in cell experiment. Therefore, the role and the mechanism of 8-oxoG formation might be different. However, there are limitations to reproduce the intrinsic oxidative stress condition in *in vitro*, and it requires some improvement of cell culture condition.

In conclusion, we demonstrated that the promoter regions of adipose tissue-specific genes are GC-rich, which could be important for the epigenetic function of the 8-oxoG.

Furthermore, genes with GC-rich transcription factor binding sites in their promoters became more active with increasing 8-oxoG abundance as demonstrated by Sp1- and ERE-luciferase assays in HEK293T cells under oxidative stress condition. These results suggest that 8-oxoG promotes transcription during adipose tissue development in mice.

## METHODS

### Tissue collection

C57BL/6 female mice were obtained from Koatech (Pyeongtaek, South Korea) and were individually housed in ventilated cages with free access to food and water. Mice (n = 5) were humanely sacrificed at 4 weeks old and liver, lung and intercapular fat samples were collected and stored at −80 °C until analyses. The study protocol was approved by the Institutional Animal Care and Use Committee of Seoul National University (SNU-170912-22) and was conducted in accordance with the approved guidelines.

### DNA hydrolysis and quantification of 8-oxoG

Genomic DNA was extracted from animal tissues using the DNeasy Blood and Tissue kit (Qiagen, Valencia, CA, USA). Before extraction, each of 100 µM desferal and butylated hydroxytoluene was added to the DNA extraction solution. The extracted DNA was enzymatically hydrolyzed to generate deoxynucleosides. Briefly, 20 µg of DNA were added to 40 µl hydrolysis buffer (100 mM NH_4_HCO_3_ [pH 7.6], 10 mM MgCl_2_, and 1 mM CaCl_2_) along with 1 U DNase I, 0.2 mU phosphodiesterase I, and 0.1 U alkaline phosphatase, followed by incubation at 37 °C for 6 h. The samples were vacuum-centrifuged at 55 °C until they were completely dried. All reagents including nucleoside standards were purchased from Sigma-Aldrich (St. Louis, MO, USA). The pellets were dissolved in 50 µl of 5% methanol and subjected to RP-LC/MS for 8-oxoG measurement. Samples were spiked with 10 nM of 8-[^15^N_5_]oxoG internal standard with 288.9/173 mass transition (Cambridge Isotope Laboratories, Inc., Tewksbury, MA, USA). Separate samples were subjected to high-performance liquid chromatography (HPLC) for dG and dC measurements as previously described (Chepelev et al. 2015).

The RP-LC/MS system consisted of an Ultimate 3000 RS HPLC instrument (Dionex, Sunnyvale, CA, USA) coupled with a TSQ ENDURA triple-quadrupole mass spectrometer (Thermo Fisher Scientific, Waltham, MA, USA). A Kinetex reversed-phase chromatography column (2.1×100 mm, inner diameter = 2.6 µm, Phenomenex, Torrance, CA, USA) was used with an injection volume of 10 µl. For separation and quantification of each nucleoside, a gradient program was established with solvent A (0.1% formic acid in water) and solvent B (0.1% formic acid in methanol), starting with 5% solvent B for 0.5 min, then ramping to 90% solvent B over 6 min, holding at 90% solvent B for 1.5 min, and re-equilibrating with 5% solvent B for 5 min at a flow rate of 300 µl/min. Mass spectrometric detection was performed using positive electrospray ionization in multiple reaction monitoring mode to monitor the 284.1/168.2 mass transition of 8-oxoG. For HPLC analysis, separation was achieved with three mobile phases - i.e., solvent A, deionized water; solvent B, 50 mM (NH_4_)_2_HPO_4_ (pH 4.0 with phosphoric acid); solvent C, methanol. The gradient program for the Shiseido column (Capcell Pak C18 UG120 4.6 × 250 mm, inner diameter = 5 µm, Phenomenex) was 82.5% solvent A, 15% solvent B, and 2.5% solvent C from 0-3 min; 55% solvent A, 15% solvent B, and 30% solvent C from 3-8 min; 82.5% solvent A, 15% solvent B, and 2.5% solvent C from 8-8.5 min; and 82.5% solvent A, 15% solvent B, and 2.5% solvent C from 8.5-10 min at a flow rate of 1.0 mL/min at 40 °C. The injection volume was 10 µl, and diode array detection was set to 280 nm.

### 8-oxoG enrichment by affinity purification

DNA fragments harboring 8-oxoG were enriched as previously described (Ding et al. 2017), with minor modifications. Briefly, 5 µg of the genomic DNA extracted from adipose or lung tissues of randomly selected mice (n = 2) were sheared with an S2 ultrasonicator (Covaris, Woburn, MA, USA) in 10 mM Tris buffer (pH 8.0) to obtain ~150-bp fragments. After sonication, the fragmented DNA was concentrated to 20 µl in 100 mM NaPi buffer (pH 8.0) using QIAquick PCR purification kit (Qiagen). A 100-µl volume of 100 mM NaPi buffer containing 20 mM amine-PEG2-biotin (Thermo Fisher Scientific) was added and the mixture was heated to 75 °C for 10 min. After thermal equilibration, 5 mM K_2_IrBr_6_ was added for 1 h for 8-oxoG biotinylation. The DNA fragments biotinylated at 8-oxoG were eluted with 125 µl Tris buffer using the QIAquick PCR purification kit and extracted using Dynabeads MyOne Streptavidin C1 (Thermo Fisher Scientific); strands complementary to those with bound biotinylated 8-oxoG were released by incubation in 150 mM NaOH at 20 °C for 30 min, and concentrated to 10 µl ddH_2_O using ssDNA/RNA Clean & Concentrator kit (Zymo Research, Irvine, CA, USA).

### 8-oxoG sequencing (OG-seq)

The 8-oxoG-enriched DNA fragments obtained by affinity purification were subjected to next-generation sequencing. Libraries were constructed using the TruSeq Nano DNA kit (Illumina, San Diego, CA, USA) and sequenced with the TruSeq SBS Kit v3-HS on a HiSeq 2000 sequencer (Illumina) to obtain 101-bp paired-end reads. Image analysis and base calling were performed using the Illumina pipeline (v1.8) with default settings. The reads were aligned to the mouse reference genome (mm10) with Isaac aligner (Illumina). Duplicated reads were identified and removed with Picard (http://broadinstitute.github.io/picard/); enriched peaks were called from the mapped reads using MACS2 (Zhang et al. 2008), and peaks were annotated with ChIPseeker (Yu et al. 2015). 8-OxoG lesions differing significantly from those in input DNA (*P* < 10^-4.5^) were selected for further analyses.

### RNA sequencing

Total RNA was extracted from adipose or lung tissue of two randomly selected mice using the RNeasy Mini kit (Qiagen) with DNase I (Qiagen) treatment. RNA integrity was evaluated with a Bioanalyzer (Agilent Technologies, Santa Clara, CA, USA). RNA sequencing libraries were generated using the TruSeq RNA sample Preparation kit (Illumina) and were sequenced on the HiSeq 2000 sequencer, which yielded ~100 million paired-end reads (2×101 bp). TopHat alignment coupled with Cufflinks assembly was used to generate the final transcriptome assembly and examine gene expression levels (Ghosh and Chan 2016). The number of reads aligned to each gene was normalized to fragments per kilobase of exon per million (FPKM).

### Functional enrichment and analysis

Genes in each tissue with promoters harboring 8-oxoG lesions were categorized as inactive for FPKM = 0 (designated as an “off” gene), moderately active for FPKM ≤ 2^9^ (designated as a “low” gene), and active for FPKM > 2^9^ (designated as a “high” gene). Biological functions and transcription factor binding motifs of the categorized genes were evaluated by Gene Set Enrichment Analysis (http://software.broadinstitute.org/gsea) (Subramanian et al. 2005). The query gene set was computationally overlapped with the Molecular Signature Database using a false discovery rate *Q*-value cutoff of 0.05. Transcription factor binding sites were predicted using AliBaba 2.1 (Grabe 2002) with the read DNA sequence obtained by 8-oxoG sequencing. The biological relevance of the genes of interest was investigated through multiplex literature mining using PubMatrix (Becker et al. 2003).

### Luciferase reporter gene assay

HEK293T cells (5×10^4^ cells per well) were seeded in a 12-well plate and cultured for 24 h before transfection with Sp1-luciferase reporter plasmid DNA (0.5 g; Panomics, Fremont, CA, USA) or a 3× ERE TATA luc construct (Addgene, Cambridge, MA, USA) for 24 h. The medium was then replaced and cells were treated with 300 µM H_2_O_2_ with or without pre-treatment of 500 µM N-acetylcysteine (NAC) for 5 min, followed by incubation for 3 h. The luciferase activity of cell lysates was measured using a luciferase assay kit (Promega, Madison, WI, USA) according to the manufacturer’s instructions. Relative luciferase activity was normalized to total protein content of the lysates.

### Immunofluorescence assay

HEK293T cells (5×10^4^ cells per well) were seeded onto coverslips in 12-well plate 24 h prior to staining. Cells were treated with 300 µM H_2_O_2_ with or without pre-treatment of 500 µM NAC for 5 min, followed by incubation for 3 h. The cells were then washed with PBS and fixed in 4% paraformaldehyde (Electron Microscopy Sciences, Hatfield, PA, USA) for 15 min at room temperature. The cells were washed twice with PBS and permeabilized with 0.2% Triton X-100 in PBS for 10 min. The cells were incubated with an antibody against 8-oxoG (Merck KGaA, Darmstadt, Germany) diluted 1:200 in 0.05% Tween 20 in PBS for overnight at 4 °C. Nuclei were stained with DAPI (Life technologies, Carlsbad, CA, USA) for 1 hr at room temperature and analyzed using an LSM 700 (Zeiss, Oberkochen, Germany).

### Cytotoxicity assay

HEK293T cells (5×10^4^ cells per well) were seeded 6-well plate 24h prior to cytotoxicity assay. Cells were treated with 300 µM H_2_O_2_ with or without pre-treatment of 500 µM NAC for 5 min, followed by incubation for 3 h. The cells were washed with PBS and collected by incubation with trypsin and the viable cells were counted using a hemocytometer after trypan blue staining (Thermo Fisher Scientific).

## DATA ACCESS

RNA-seq and OG-seq raw and processed data from this study have been submitted to the NCBI Gene Expression Omnibus (GEO; http://www.ncbi.nlm.nih.gov/geo/) under accession number GSE124359 and GSE124712.

## ACKNOWLEDGEMENTS

Author contributions: J.W.P., Y.I.H., S.C.Y., and T.M.K. conceived the study, designed and performed experiments, analysed and interpreted the data, and wrote the manuscript. J.P. and J.K. conceived the study, designed the experiments, interpreted the data, performed bioinformatics analysis, and wrote the manuscript.

## FUNDING

This work was supported by the Research Resettlement Fund for the new faculty of Seoul National University (1403-0160066 to J.P.) and the Basic Science Research Program through the National Research Foundation, the Ministry of Education, Korea (NRF-2017R1A6A1A03015713 to J.K.).

## DISCLOSURE DECLARATION

The authors declare no conflict of interest.

