## Supplementary figures and images for "8-OxoG in GC-rich Sp1 binding sites enhances gene transcription during adipose tissue development in juvenile mice"

### Supplementary Figure 1

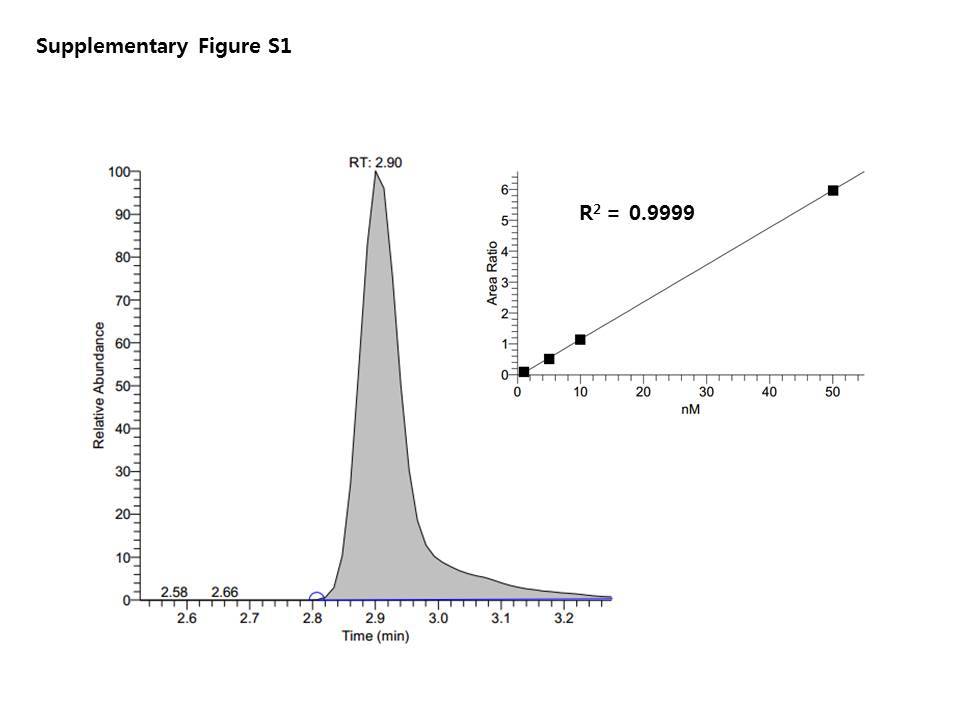

### Supplementary Figure 2

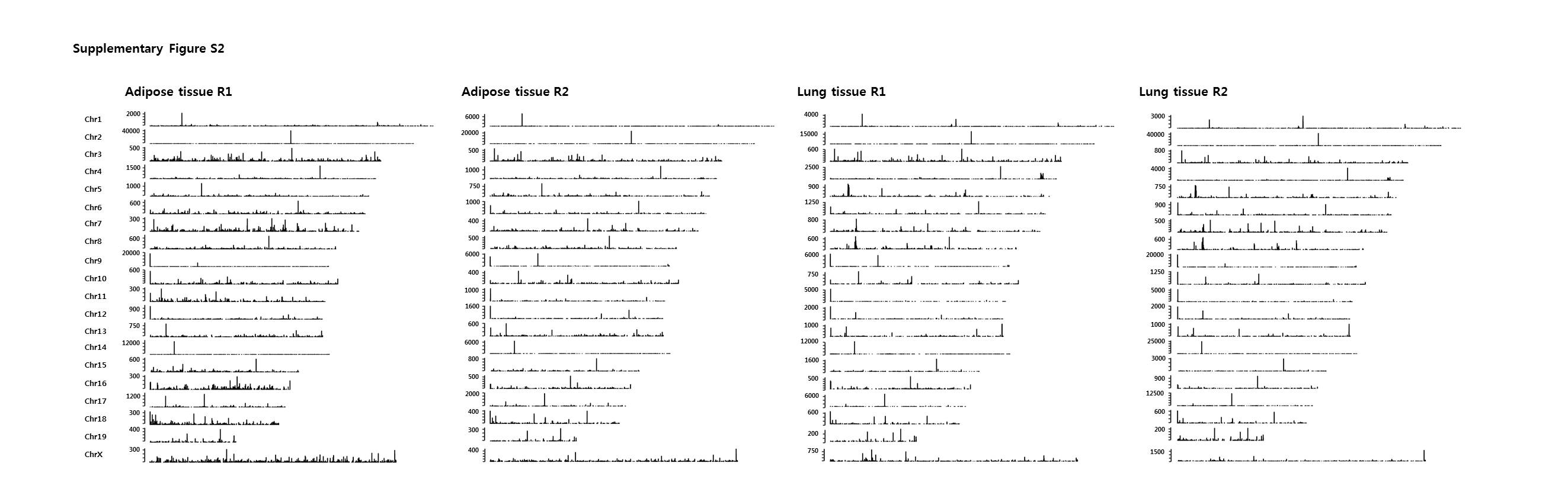

### Supplementary Figure 3

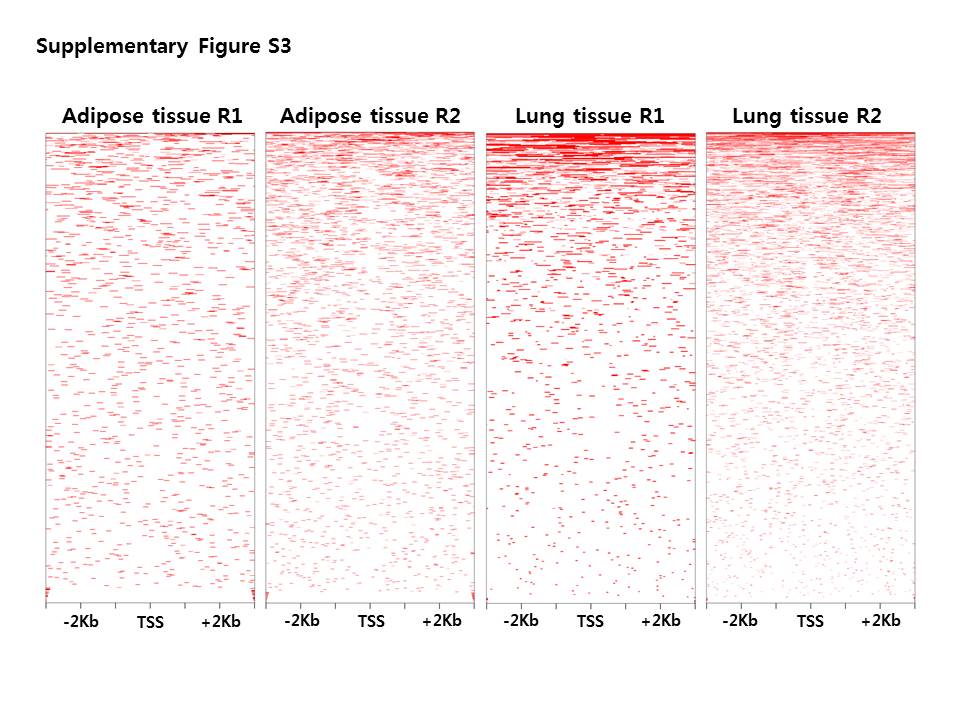

### Supplementary Figure 4

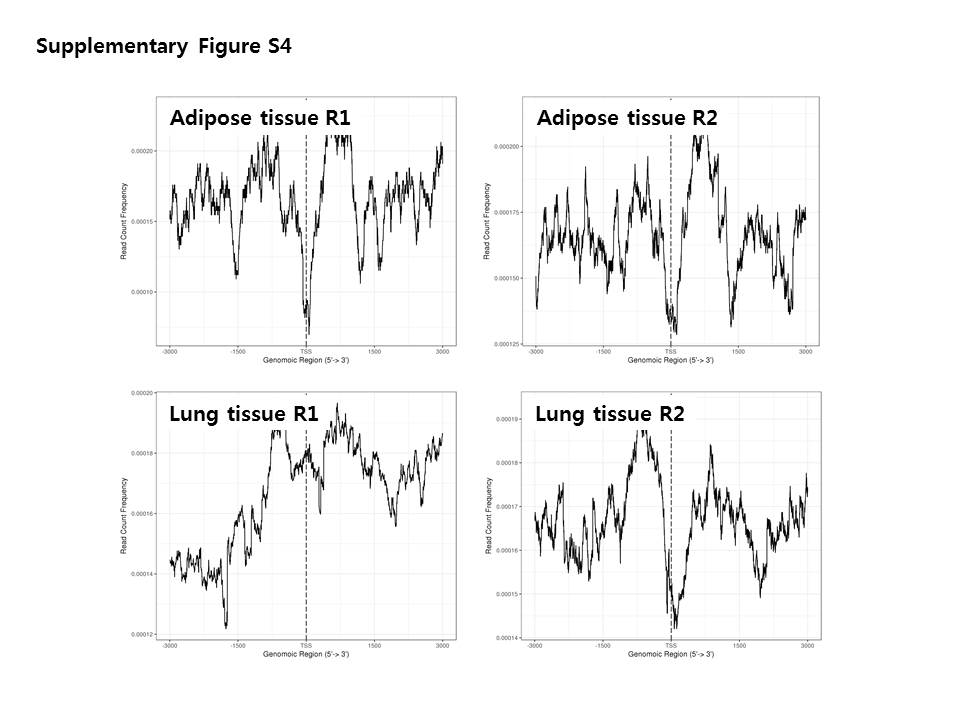

### Supplementary Figure 5

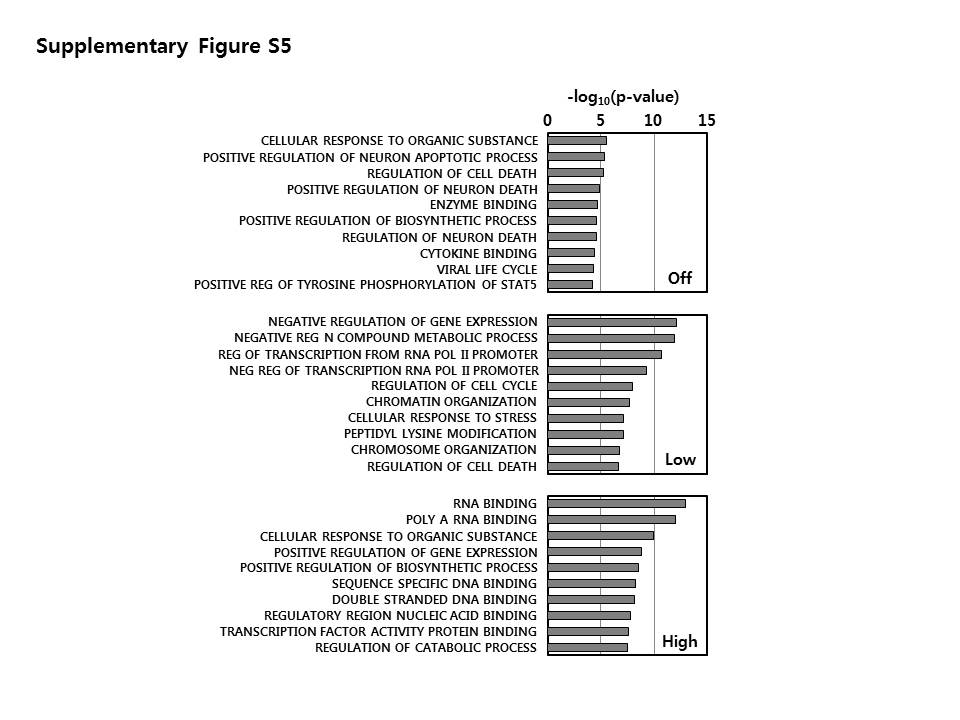

### Supplementary Figure 6

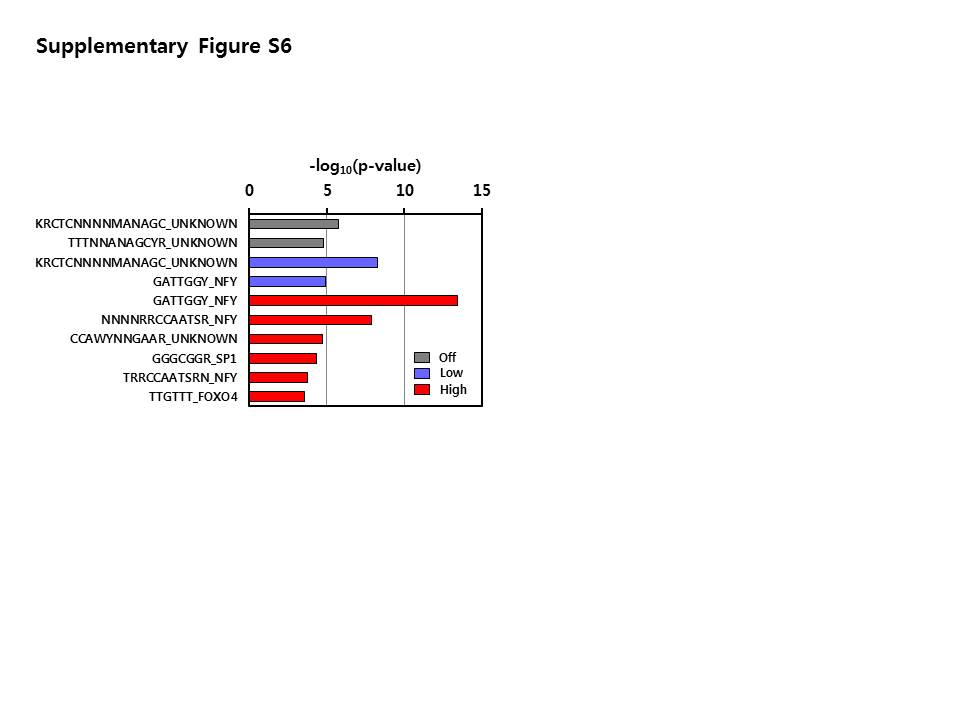
